# Genetic reduction of the translational repressors FMRP and 4E-BP2 preserves memory in mouse models of Alzheimer’s disease

**DOI:** 10.1101/2024.10.08.617315

**Authors:** Felipe C. Ribeiro, Danielle Cozachenco, Jean-Claude Lacaille, Karim Nader, Fernanda G. De Felice, Mychael V. Lourenco, Argel Aguilar-Valles, Nahum Sonenberg, Sergio T. Ferreira

**Author notes:** These authors contributed equally to this work (Felipe C. Ribeiro, Danielle Cozachenco). Corresponding authors: Sergio T. Ferreira, Nahum Sonenberg and Argel Aguilar-Valles.

## Abstract

Alzheimer’s disease (AD) is characterized by progressive memory decline. Converging evidence indicates that hippocampal mRNA translation (protein synthesis) is defective in AD. Here, we show that genetic reduction of the translational repressors, Fragile X messenger ribonucleoprotein (FMRP) or eukaryotic initiation factor 4E (eIF4E)-binding protein 2 (4E-BP2), prevented the attenuation of hippocampal protein synthesis and memory impairment induced by AD-linked amyloid-β oligomers (AβOs) in mice. Moreover, genetic reduction of 4E-BP2 rescued memory deficits in aged APPswe/PS1dE9 (APP/PS1) transgenic mouse model of AD. Our findings demonstrate that strategies targeting repressors of mRNA translation correct hippocampal protein synthesis and memory deficits in AD models. Results suggest that modulating pathways controlling brain mRNA translation may confer memory benefits in AD.

## Main

Alzheimer’s disease (AD) is the most prevalent cause of dementia in the elderly worldwide^1^ and is characterized by progressive cognitive decline^2^. Hippocampal mRNA translation (protein synthesis) is crucial for memory consolidation by promoting synapse strengthening^3^. Signaling pathways controlling the initiation and elongation steps of mRNA translation are deregulated in the brains of AD patients and mouse models^4,5^. This results in attenuation of global brain protein synthesis, changes in brain proteome composition^6^ and memory failure in AD mouse models^4,7^. These findings suggest that multiple pathways converge to arrest hippocampal translation, thereby contributing to AD pathogenesis.

Fragile X messenger ribonucleoprotein (FMRP, encoded by the *Fmr1* gene) is a negative regulator of protein synthesis that blocks both initiation and elongation phases of mRNA translation^8^, but it is unknown whether FMRP plays a role in AD^9^. Eukaryotic initiation factor 4E-binding protein 2 (4E-BP2, encoded by the *Eif4ebp2* gene) is the major 4E-BP isoform in the brain^10^. 4E-BP2 is a translational repressor that binds to eIF4E and prevents formation of the eIF4F complex^11^. Here, we tested the hypothesis that genetic reduction of FMRP or 4E-BP2 could rescue memory deficits in AD mouse models.

We initially investigated whether genetic reduction of FMRP or 4E-BP2 prevented the attenuation of hippocampal protein synthesis induced by amyloid-β oligomers (AβOs), soluble neurotoxins that cause synapse failure and cognitive impairment in the AD brain^2,7^. We infused 10 pmol AβOs (or vehicle) via intracerebroventricular (i.c.v.) injections in wild type (WT), *Fmr1*^y/-^ or *Eif4ebp2*^+/-^ mice, and assessed global *de novo* protein synthesis using surface sensing of translation (SUnSET) in *ex vivo* hippocampal slices. In line with our previous reports^5,12–14^, we found that i.c.v. infusion of AβOs caused a reduction of approximately 30% in *de novo* protein synthesis in hippocampal slices obtained from WT mice **(Figure 1A, B; Supplementary File 1; Supplementary Table 1)**. In contrast, infusion of AβOs did not attenuate hippocampal protein synthesis (compared to vehicle-infused mice) in either *Fmr1*^y/-^ or *Eif4ebp2*^+/-^ mice **(Figure 1A, B)**.

**Figure 1.**
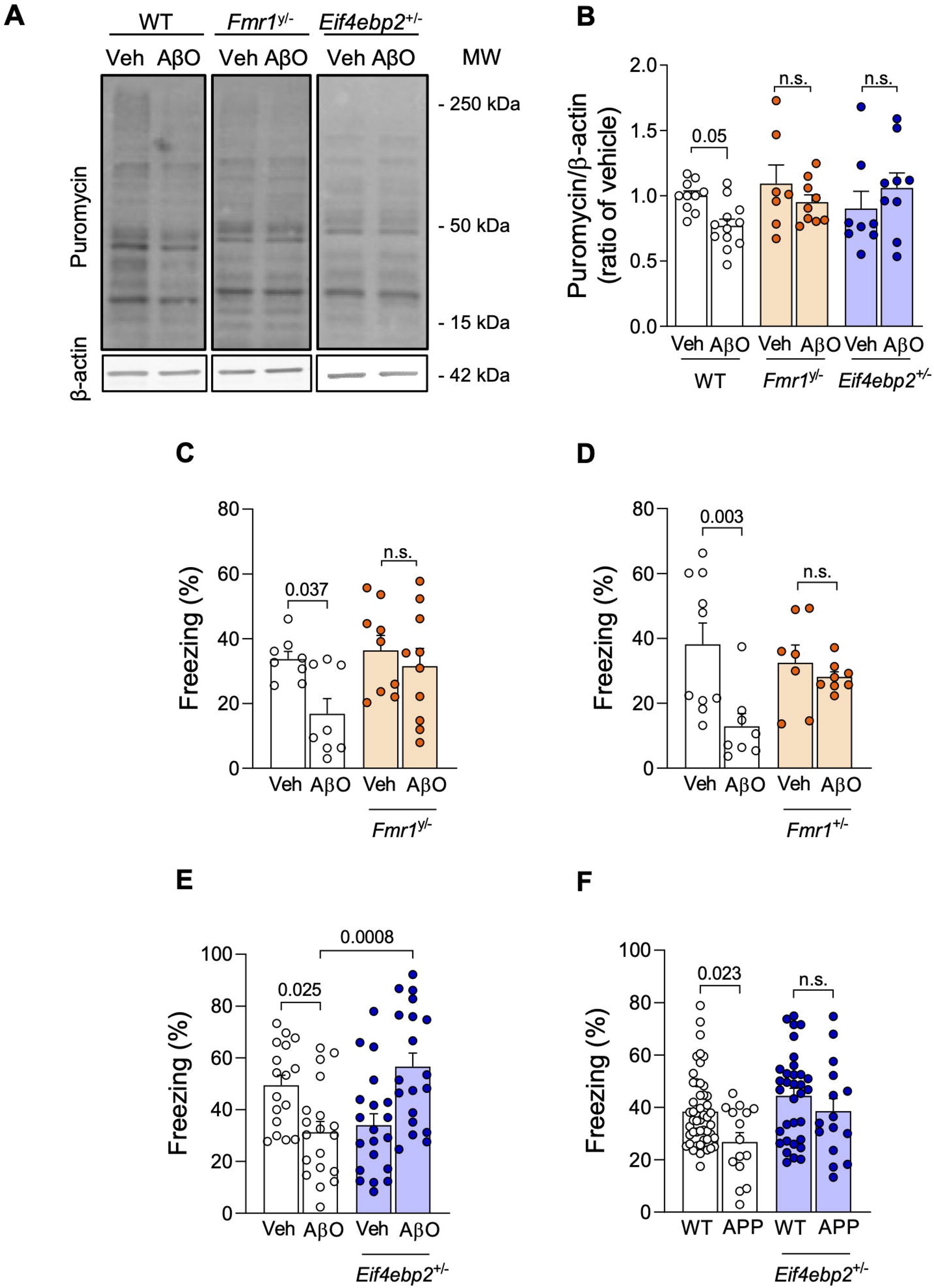
Genetic reduction of FMRP or eIF4E-BP2 rescues defective hippocampal protein synthesis and memory in AβO-infused mice. (A, B) 3-month-old male *Eif4ebp2*^+/-^ or *Fmr1*^y/-^ mice (or corresponding WT littermates) received an i.c.v. infusion of 10 pmol AβOs (or vehicle) (N = 10 Veh, 12 AβOs, 8 Fmr1^y/-^ Veh, 9 Fmr1^y/-^ AβOs, 7 Eif4ebp2^+/-^ Veh, 9 Eif4ebp2^+/-^ AβOs). Hippocampal slices were obtained 7 days later and were exposed to puromycin (5 μg/mL; 45 min) for SUnSET. Puromycin levels were evaluated by Western blotting and normalized by β-actin levels. (C-E) 3 month-old *Fmr1*^y/-^ (N = 8 Veh, 9 AβOs, 9 Fmr1^/-^ Veh, 10 Fmr1^y/-^ AβOs), *Fmr1*^+/-^ (N = 10 Veh, 8 AβOs, 7 *Fmr1*^+/-^ Veh, 8 *Fmr1*^+/-^ AβOs) or *Eif4ebp2*^+/-^ mice (N = 17 Veh, 19 AβOs, 20 *Eif4ebp2*^+/-^ Veh, 19 *Eif4ebp2*^+/-^ AβOs) (or corresponding WT littermates) received an i.c.v. infusion of 10 pmol AβOs (or vehicle) and memory was assessed using the contextual fear conditioning (CFC) test 7 days later. (F) 12-month-old APPswe/PS1dE9/*Eif4ebp2*^+/-^ mice and corresponding controls (APPswe/PS1DE9/*Eif4ebp2*^*+/+*^ mice) were assessed in the CFC memory test (N= 49 WT, 15 APP/PS1, 33 WT *Eif4ebp2*^*+/-*^, 15 APP/PS1 *Eif4ebp2*^*+/-*^); p-values are shown for appropriate comparisons; Two-way ANOVA followed by Holm-Sidak post-hoc test.

Next, we assessed the impact of AβOs on hippocampus-dependent fear memory using a contextual fear conditioning paradigm in *Fmr1*^y/-^ males, *Fmr1*^+/-^ females or WT mice. Consistent with our previous reports^7,12^, i.c.v. infusion of AβOs blocked long-term fear memory in WT mice **(Figure 1C, D; Supplementary Table 2)**. Remarkably, *Fmr1*^y/-^ or *Fmr1*^*+/-*^ mice that received infusions of AβOs exhibited intact memory in the fear conditioning test **(Figure 1C, D)**.

Next, *Eif4ebp2*^+/-^ or WT mice were tested in the contextual fear conditioning paradigm 7 days after a single i.c.v infusion of 10 pmol AβOs (or vehicle). Consistent with the notion that translational control by 4E-BP2 plays an essential role in memory processes and must be tightly regulated^10^, 4E-BP2 haploinsufficiency *per se* led to memory impairment in vehicle-infused mice **(Figure 1E)**. However, 4E-BP2 haploinsufficiency prevented AβO-induced impairment of fear memory **(Figure 1E)**. Altogether, these findings indicate that alleviating translational repression via genetic reduction of 4E-BP2 or FMRP prevents AβO-induced memory impairments.

To determine whether alleviating translational repression could represent an effective approach to rescue memory in a chronic AD model, we generated APPswe/PS1dE9 (APP/PS1) mice harboring a single 4E-BP2 allele (*APP/PS1/4E-BP2*^*+/-*^ mice). APP/PS1 mice develop age-related Aβ accumulation and memory impairment^12^. Whereas aged APP/PS1 mice (harboring two copies of the *4E-BP2* allele) exhibited impaired long-term fear memory, *4E-BP2* haploinsufficient APP/PS1 mice exhibited memory performance comparable to that of wildtype mice **(Figure 1F)**.

Our results demonstrate that genetic brake-release of hippocampal mRNA translation preserves memory in both acute and chronic mouse models of AD. The findings using mice with genetically reduced 4E-BP2 or FMRP support the concept that correcting brain protein synthesis defects counteracts AD-linked memory impairment. Interestingly, we did not observe significant differences in baseline hippocampal protein synthesis in *Fmr1*^y/-^ or *Eif4ebp2*^+/-^ mice compared to WT mice. Thus, while genetic reduction of FMRP or 4E-BP2 did not appear to have a major impact on global hippocampal protein synthesis, it was sufficient to block the attenuation of protein synthesis induced by AβOs. Of note, while the brain proteome of *Fmr1* KO mice has been extensively investigated (as reviewed in^15^), including changes in activity-dependent synaptic proteins, little is known on how deficiency of 4E-BP2 alters protein composition in the brain. Thus, the detailed mechanisms by which genetic reduction of *Fmr1* and *4E-BP* are protective remain elusive and further studies are warranted to elucidate them.

In conclusion, while we and others have previously demonstrated that defective proteostasis regulation^13,16^ and translational control^4,5,7^ play important roles in AD, the current study underscores the importance of two key mediators of translation in Aβ-induced memory impairments. Our findings warrant future mechanistic investigation into how these multiple dysregulated pathways converge to trigger AD-linked symptoms. The notion that impaired control of mRNA translation may be a druggable target may pave new roads toward effective therapeutics in AD.

## Methods

### Animals

Adult male or female mice (P90) were obtained from the animal facility at McGill University and were housed in groups of 2-5 mice per cage, with food and water *ad libitum*, on a 12 h light/dark cycle. Male and female APPswe/PS1dE9 mice (Jackson Laboratories, strain #005864), *Eif4ebp2*^+/-^, *Fmr1*^+/-^ and *Fmr1*^y/-^ mice on a C57BL/6J background (obtained from The Jackson Laboratories) were bred at our facilities. All procedures followed the Canadian Council on Animal Care guidelines and were approved by McGill University Committees.

### AβO preparation and intracerebroventricular (i.c.v.) infusion

Oligomerization of the Aβ_1-42_ peptide (California Peptide, Salt Lake, CA) and intracerebroventricular (i.c.v.) infusion of 10 pmol AβOs (or vehicle) in mice were performed as described^7,17^

### Surface sensing of translation (SUnSET) in hippocampal slices

Hippocampal slices (400 μm) were obtained and kept in artificial cerebrospinal fluid (aCSF; 124 mM NaCl, 4.4 mM KCl, 1 mM Na_2_HPO_4_, 25 mM NaHCO_3_, 2 mM MgCl_2_, 2mM CaCl_2_, 10 mM glucose) for 2 h. Newly synthesized polypeptides were detected using SUnSET as described^5^. Hippocampal slices were prepared 7 days after i.c.v. infusions of 10 pmol AβOs (or vehicle) in *Eif4ebp2*^+/-^, *Fmr1*^y/-^ or corresponding littermate WT mice. Slices were exposed to puromycin (5 μg/mL; 45 min) and collected for Western blotting.

### Western blotting

Hippocampal slices were dissociated using Bio-Plex Cell Lysis Kit (Bio-Rad), centrifuged for 10 min at 10,000 g at 4°C, and the supernatant was collected^13^. Protein concentration was determined using Bradford Protein Assay Kit. 50 μg total protein were resolved in 12% Tris-glycine SDS-PAGE gels, transferred to nitrocellulose membranes, and incubated with anti-puromycin (12D10 clone; 1:1,000; EMD Millipore) and β–actin (1:10,000; Abcam). Immunoblots were developed using IR dye-conjugated fluorescent secondary antibodies (1:5,000; LiCor), imaged on an Odyssey system, and analyzed using ImageJ (NIH).

### Contextual fear conditioning

In the training phase, mice were placed in the apparatus and received two foot shocks (0.35 mA for 1 s at 2 min and 2:30 min), and were placed back in their home cages 30 s later. For the test phase, 24 h later, mice were placed in the same apparatus for 5 min, with no shock. Freezing behavior was recorded automatically on Freeze Frame (Harvard Apparatus, Holliston, MA).

### Statistical analysis

Data are expressed as means ± S.E.M. and were analyzed using GraphPad Prism 8 software (La Jolla, CA). Sample sizes, statistical tests, and p-values for each experiment are indicated in the figure legend.

## List of abbreviations

4E-BP2: eukaryotic initiation factor 4E (eIF4E)-binding protein 2;
AβOs: amyloid-β oligomers
AD: Alzheimer’s disease
CYFIP1: cytoplasmic FMR1-interacting protein 1
eIF2α: eukaryotic translation initiation factor 2α
eIF4E: eukaryotic translation initiation factor 4E
eEF2: eukaryotic elongation factor 2
FMRP: Fragile X messenger ribonucleoprotein
ISR: integrated stress response
ISRIB: integrated stress response inhibitor
SUnSET: surface sensing of translation
WT: wild type

## Declarations

### Ethics approval and consent to participate

All procedures followed the Canadian Council on Animal Care guidelines and were approved by McGill University Committees.

### Consent for publication

Not applicable.

### Availability of data and materials

The datasets used and/or analyzed during the current study are available from the corresponding author upon reasonable request.

### Author contributions

F.C.R., D.C., A.A.-.V. and S.T.F. designed the study. F.C.R., D.C., and A.A.-.V. performed research. F.C.R., D.C., and A.A.-.V. analyzed data. K.N., M.V.L., J.C.-.L., F.G.D.F., A.A.-.V., N.S. and S.T.F. contributed reagents, materials, animals and analysis tools. F.C.R., D.C., F.G.D.F., M.V.L., A.A.-.V., N.S. and S.T.F. analyzed and discussed results. F.C.R., D.C., M.V.L. and S.T.F. wrote the manuscript with inputs from the other authors.

## Acknowledgements

Not applicable.

