## Supplementary figures and images for "Genetic reduction of the translational repressors FMRP and 4E-BP2 preserves memory in mouse models of Alzheimer’s disease"

### Supplementary file 1.pdf

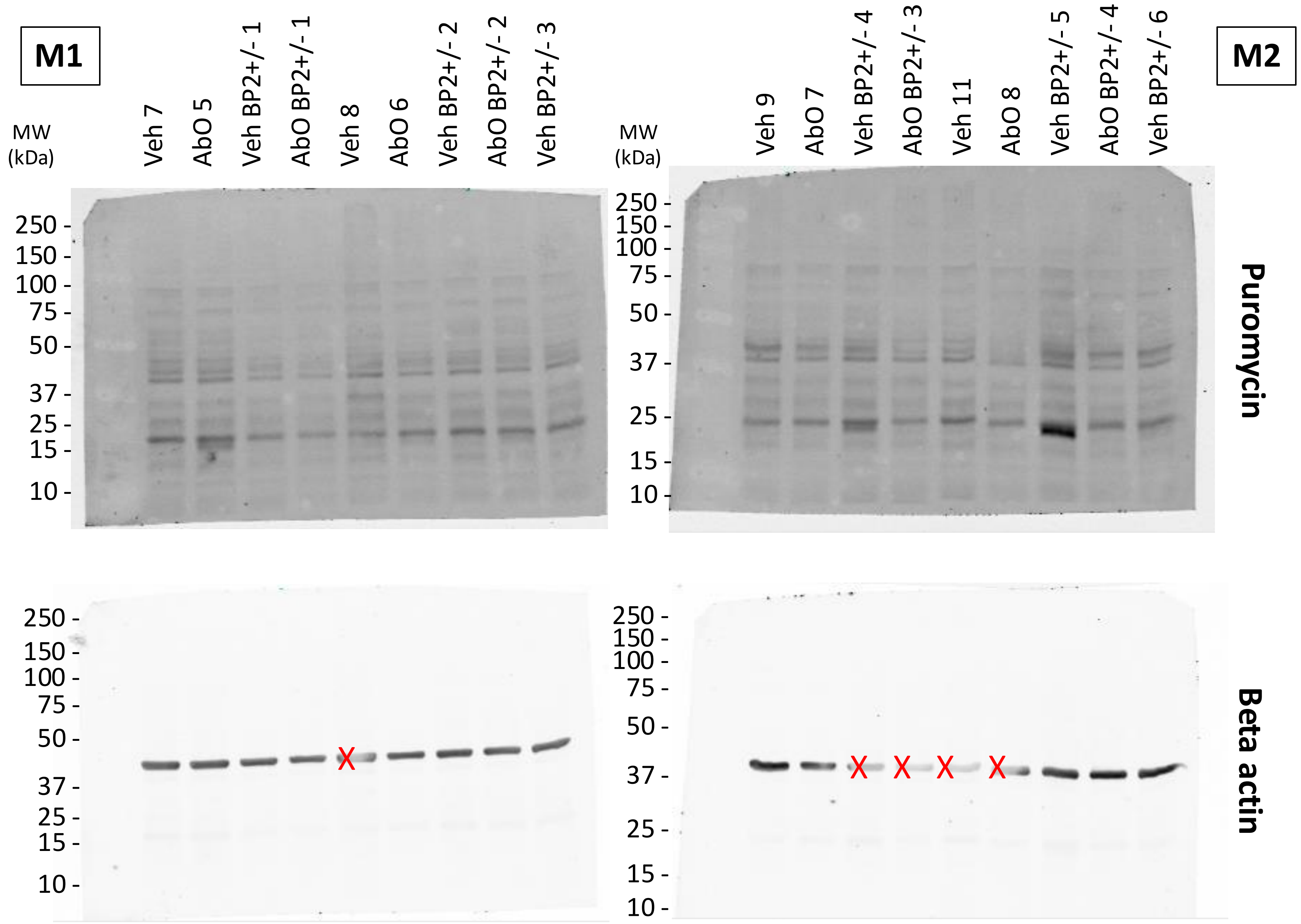


Ribeiro et al.

X – band too clear – we discarded

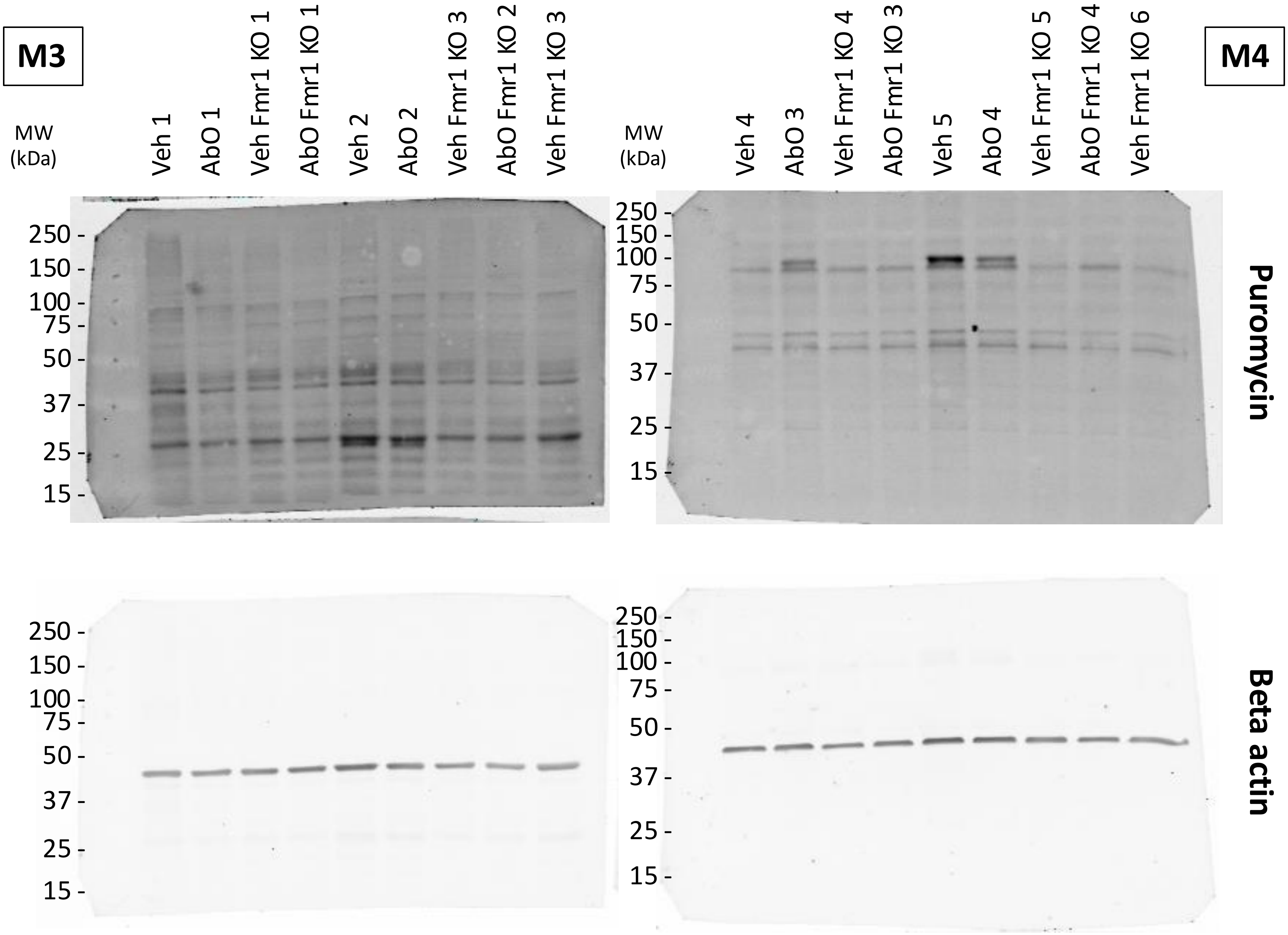


Ribeiro et al.

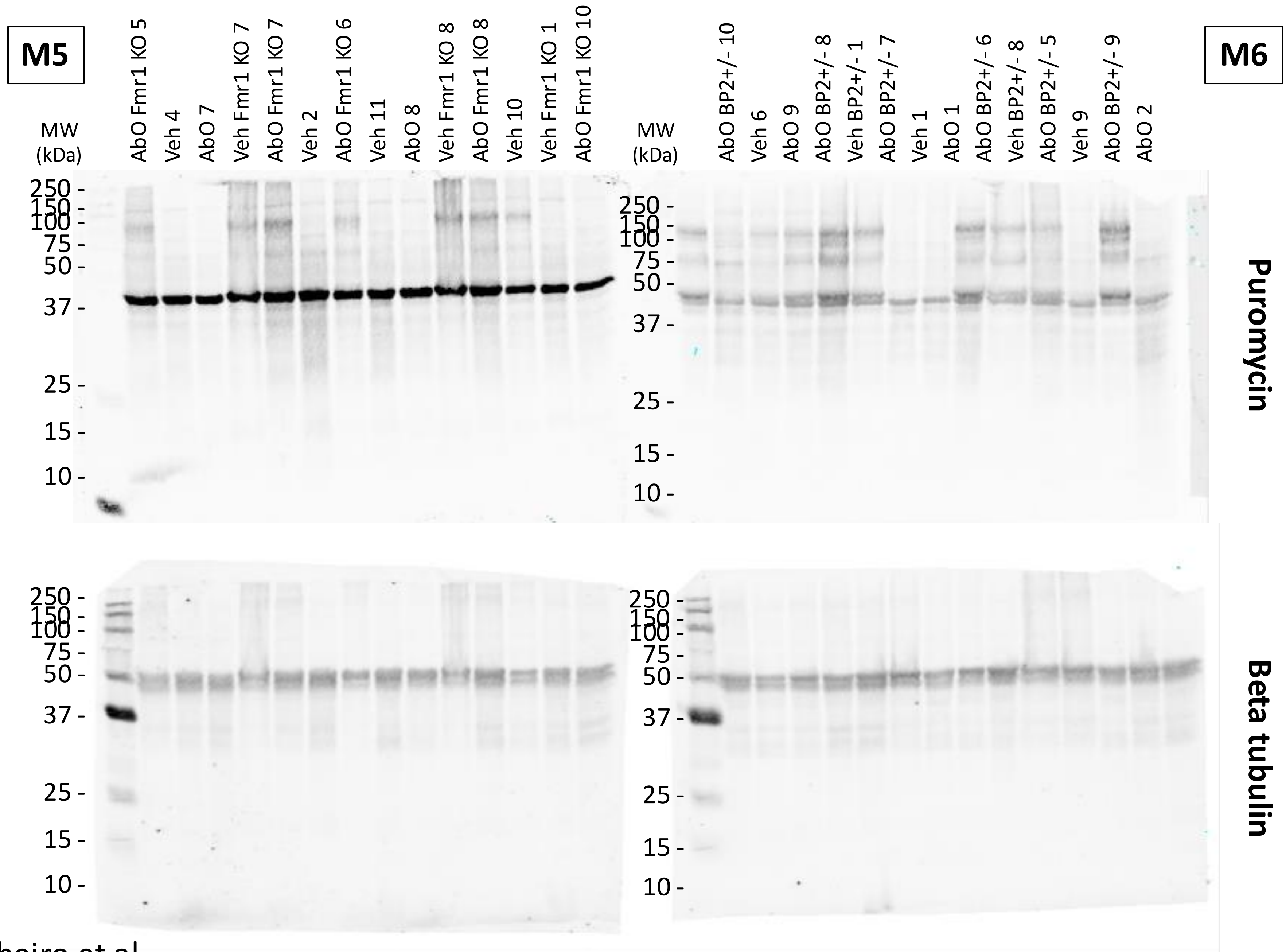


Ribeiro et al.
